# UP-REGULATION OF SYNAPTOBREVIN-2 TO DELAY AGE-RELATED COGNITIVE IMPAIRMENT

**DOI:** 10.64898/2026.05.05.722770

**Authors:** Jacob Bradford Miller, Ankit Seth, Ashiq Mahmood Rafiq, Weiping Han, Ferenc Deak

## Abstract

The slowing of executive function and memory impairment are the leading hallmarks of cognitive decline with age. The exact cause of this change is unknown and is the focus of aging research. Expression levels of Vesicle Associated Membrane Protein 2 (VAMP2)/Synaptobrevin-2 (syb2) are decreased with age. Here we report results from a novel transgenic mouse model (TgV2) that overexpresses syb2. We hypothesized that overexpression of syb2 improves synaptic function during aging, thus it delays dementia. Aged TgV2 mice, which maintained syb2 levels, performed better in spatial memory tests than 2-year-old WT control mice, which had lost half of syb2 due aging. In hippocampal CA1 synapses of aged TgV2 mice, long-term potentiation was increased. These effects of maintained syb2 levels were beneficial for both males and females providing improved synaptic plasticity. These results indicate that overexpression of syb2 supports cognitive function throughout the aging process and better resist age-related synaptic dysfunction.

## Introduction

Age-related cognitive decline is associated with deficits in executive functions such as visuo-spatial working memory and cognitive flexibility, in addition to long-term episodic memory and spatial orientation ^1-3^. While dementias such as Alzheimer’s disease (AD) have distinctive pathological markers like tau fibrillary tangles and amyloid beta plaques^4^, as well as significant neuronal loss linked to cognitive decline, sources of cognitive decline in the normal aging population remain elusive. Although multiple factors were associated with brain aging and cognitive decline, proof of their causative role in dementia is lacking. Additional and alternative pathological hallmarks have been previously attributed to age-related memory loss including decrease in synaptic density and strength^5-9^. Syb2 levels have shown importance in long-term studies on human patients, as it was found that higher levels of syb2 correlated with the reduced likelihood of dementia and greater cognitive function^10, 11^. Importantly, a recent study revealed that elderly patients with the highest syb2 levels are best protected from dementia^10^. As we previously reported, selective reduction of syb2 levels causes cognitive impairments ^12^. We have discovered that there are spatial learning and memory deficits in syb2 heterozygous knockouts (syb2+/-) even at young adult age, which proves that reduced syb2 levels lead to cognitive impairment. Here we present results from our next study where we have examined whether maintaining syb2 levels overcomes age related cognitive decline.

The SNARE (Soluble N-ethylmaleimide-sensitive factor attachment protein receptor) complex is necessary for proper synaptic transmission as interactions between v-snares and t-snares within the complex facilitate vesicular release at the presynaptic terminal. Vesicular Associated Membrane Protein 2 (VAMP2), also called Synaptobrevin-2 (syb2), is a vesicular SNARE protein involved in the fusion of vesicles to the synaptic membrane for exocytosis and endocytosis^13-21^. Synaptobrevin-2 is known to be evolutionarily conserved among a wide range of both vertebrates and invertebrates. For instance, human syb2 amino acids 1-94 share 99% of its identity with mouse syb2 ^22^ and a complete match with bovine syb2. As previously reported, syb2 levels significantly decrease with age and selective reduction of syb2 levels causes cognitive impairments ^23^. Briefly, knocking out syb2 in a heterozygous mouse model resulted in signs of age-related cognitive decline, with aged heterozygous mice performing worse in spatial learning and memory tasks, reduced LTP in the CA1 region of the hippocampus, and impaired vesicular release rates in hippocampal neurons ^23^. This has brought us to pose the question if using a model of syb2 overexpression would prevent memory deficits in aging.

## Results

A novel mouse model to counter brain aging, known as TgV2 ^24, 25^, was designed by using a mouse Thy1 gene promotor to express a specific fusion protein consisting of syb2 with N-terminally fused ECFP, a blue fluorescence protein. In this study, we have examined the effects of this transgene on cognitive functions alongside naturally expressed syb2 and how changes in protein levels over the aging process effects synaptic transmission. We used a variety of techniques including behavioral studies, electrophysiology and live fluorescence imaging to assess changes in spatial learning and memory and synaptic plasticity in TgV2 mice during the aging process.

### Syb2 Overexpression Benefits Long Term Spatial Memory in Aged Mice

Young and adult TgV2 mice appeared similar in behavior to WT littermates. Age-related changes were assessed using a variety of assays, including open field and rotarod tests for motor coordination. No significant difference was noted between young WT and TgV2 mice in mobility, motor coordination and anxiety (Suppl. Fig 1.). Aged mice covered less distance within 10min in both WT and TgV2 cohorts than young mice: (Young adult WT 26.56 meters vs Aged WT 14.79 meters, *p=0.0422; Tukey’s multiple comparison test), (Young adult TgV2 28.46 meters vs Aged TgV2 15.71 meters, *p=0.0135). Aged mice moved slower similarly in both WT and TgV2 cohorts than young mice (Young adult WT 44.26 mm/s vs Aged WT 24.64 mm/s, *p=0.0422; Tukey’s multiple comparison test), (Young adult TgV2 47.41 mm/s vs Aged TgV2 26.27 mm/s, *p=0.0143). The only difference noted is that TgV2 mice overexpressing syb2 tend to show less reduction by age in time being mobile in the open field box (70.96 sec immobile vs 157.57 sec for aged TgV2, p=0.1377), while immobility increased significantly for aged WT mice (68.16 Sec vs 190.33 sec; *p=0.0327).

A western blotting assay was performed using hippocampal samples which proved that TgV2 mouse line maintains high levels of the syb2 fusion protein at two years or older age **(Fig.1A)**, which is comparable to syb2 levels in young adult WT brains. Moreover, syb2/syb2-eCFP fusion protein levels are 1.8-2 folds of the natural syb2 levels compared to age matched WT brains **(Fig.1B**), which suffer from a 50% reduction due to aging.

**Figure 1:**
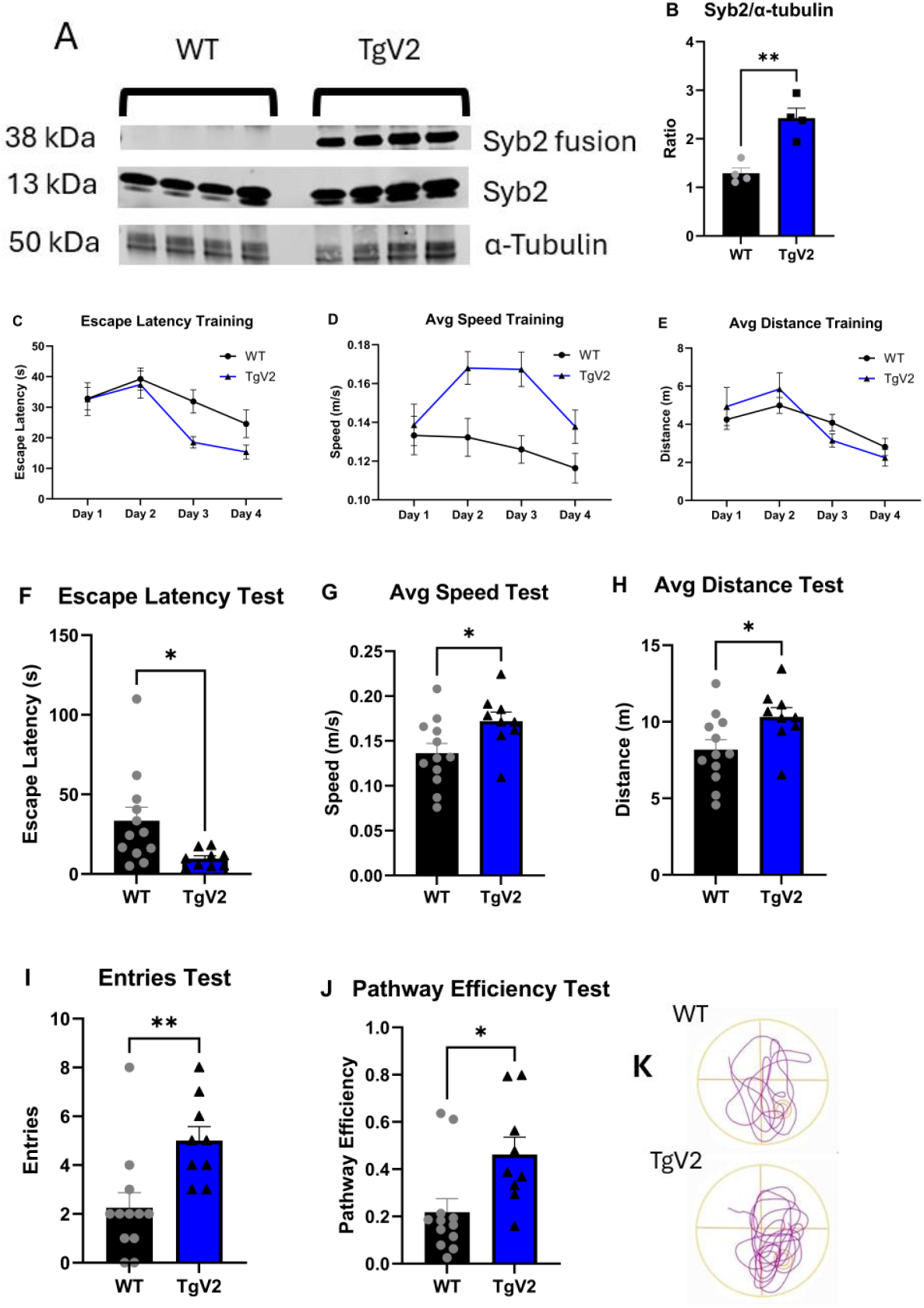
Western Blot Analysis of Syb2 in TgV2 and Tg2576 mice. **(A)** In this western plot, hippocampal samples were taken from the half brains of severely aged mice (∼2-year-old) and analyzed to prove that this fusion protein is still prevalent in old age. **(B)** Total endogenous Syb2/Syb2-eCFP fusion protein levels are 2-fold of the natural Syb2 levels in WT brains (**p=0.0029; unpaired t-test). Overexpressed syb2 protein is a fusion protein tagged with ECFP; therefore, it will not show up in the same molecular weight as standard syb2 expression. **TgV2 mice show greater spatial learning and memory during in old age.** 2-year-old mice (WT n = 12) (TgV2 n = 9) **(C)** Latency to reach the platform during 4-day training period. **(D)** Average swim speed during training. **(E)** Mice traveled less before finding the platform indicating learning. **(F)** Escape latency on probe day trial (**p=0.0227; unpaired t-test). **(G)** TgV2 mice swam faster during the probe day trial (**p=0.0325). **(H)** Distance traveled until reaching the platform zone in the probe trial (p=0.0329; unpaired t-test). **(I)** Number of entries into the platform zone (**p=0.0052). **(J)** Pathway efficiency until first entry to the platform zone (**p=0.0149). **(K)** Representative track plot images during the probe trial.

To examine the effects of Syb2 overexpression on spatial learning and memory during the aging process, we tested both young adult mice (6-8 months old) and aged mice (2-year-old) using the Morris Water Maze assay. Young adult mice showed no differences in learning and memory **(data not shown)** compared to WT mice. Although no significant differences were observed in the young adult cohorts, TgV2 mice showed a trend on test day for faster escape latency to reach the platform zone **(Fig.S1D)** and a shorter total distance traveled **(data not shown)** compared to WT Mice. For the 2-year-old cohort, it took markedly longer for WT mice to find the platform, and their search pattern was less efficient compared to young WT mice. Importantly, aged TgV2 maintained the capacity for spatial learning and memory **(Fig.1)**. TgV2 mice in the aged cohort displayed a greater capacity for long-term memory as seen during the test of recall on the location seven days later compared to WT mice. This is indicated by their better performance in multiple parameters, including escape latency, distance to first entry, number of entries to the platform zone and pathway efficiency to the first entry into the platform zone **(Fig.1F, H-J)**. We also observed that aged TgV2 mice maintain a significantly faster average swim speed compared to aged WT mice **(Fig.1G)**.

#### Higher Synaptobrevin-2 Levels Cause Improved Synaptic Plasticity

Previous data showed that syb2 deficient heterozygous knockout mice had impaired Long-Term Potentiation (LTP) in the CA1 region of the hippocampus^23^. To assess if syb2 overexpression will result in the opposite effect, we performed LTP assay on hippocampal slices from aged 18-month-old to 2-year-old mice. During the last 10 minutes of recording, enhanced maintenance of synaptic plasticity was observed in TgV2 hippocampal slices (WT = 150.8% vs TgV2 = 170.6% average normalized fEPSP) **(Fig. 2C)**. To further examine the differences in synaptic plasticity between genotypes, we recorded the input-output responses before and after the baseline recordings using a stimulus range of 0-100 µA. Output curves indicated increased amplitude and slope of the EPSP one hour after high frequency stimulation in TgV2 hippocampal slices compared to WT **(Fig.2D-E)**. These results confirmed that TgV2 mice have greater synaptic plasticity in the CA1 region of the hippocampus, in good agreement of the spatial memory improvement detected in aged TgV2 mice.

**Figure 2:**
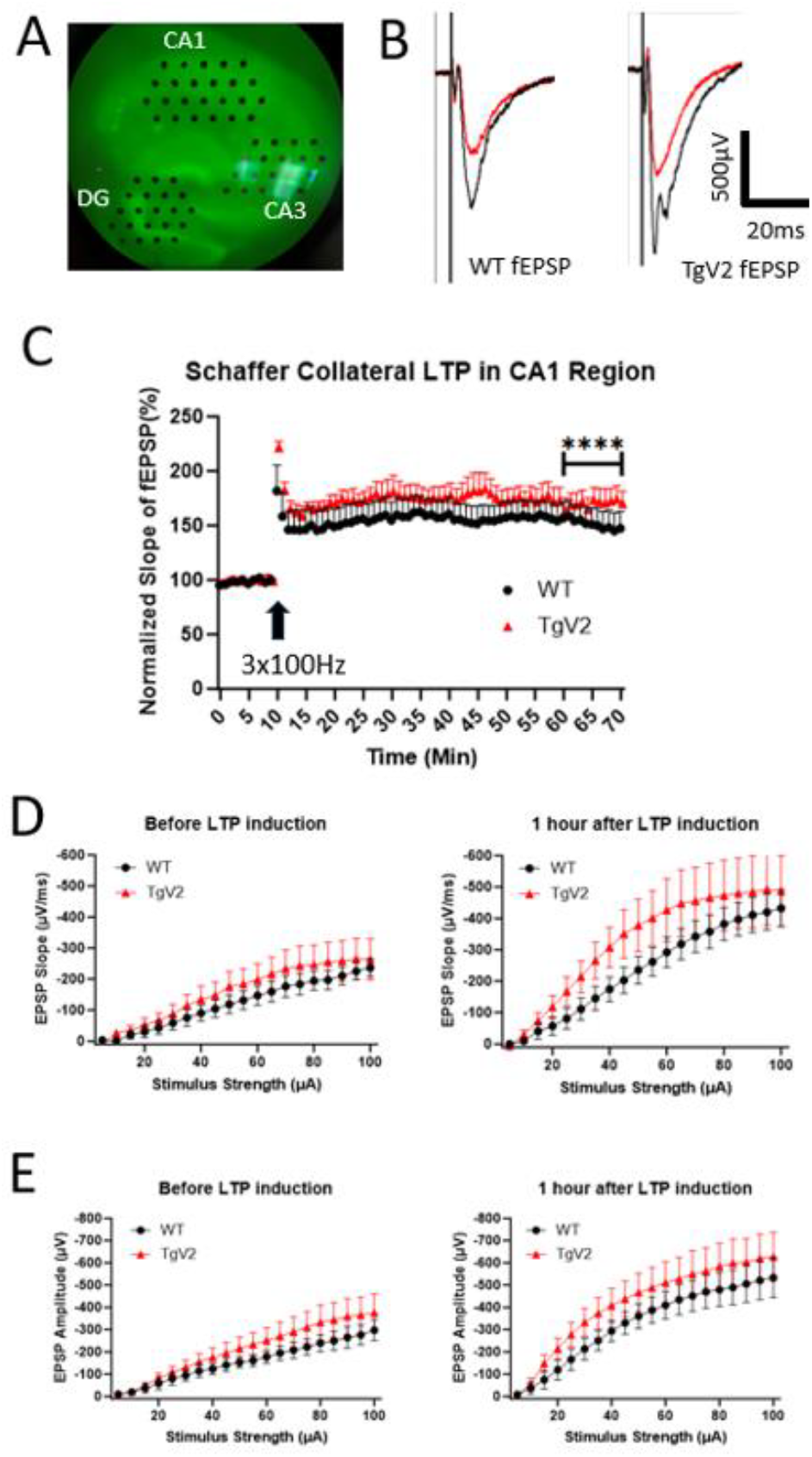
TgV2 mice have enhanced synaptic plasticity in the CA1 region of the hippocampus. (**A**) Image of hippocampal slices in microelectrode array (MEA). **(B)** Representative sample traces of fEPSP amplitude before and after LTP. **(C)** Normalized slope of fEPSP. TgV2 response was significantly higher than WT during the last 10 minutes of recording (****p<0.0001; unpaired t-test). **(D)** EPSP slope before 3×100Hz stimulation (n.s. p=0.1457; Tukey’s multiple comparisons test) and one hour after (****p<0.0001). **(E)** EPSP amplitude before stimulation (*p=0.0342) and one hour after (****p<0.0001). (WT n=4 mice, n=5 slices) (TgV2 n=5 mice, n=7 slices).∼18-24-month-old mice

#### Overexpression of Syb2 Increases vesicular release rates and Functional Synaptic Vesicles

To further observe the effects our syb2 overexpression model has on synaptic transmission, we also utilized live fluorescence imaging and whole-cell patch clamp techniques on primary cortical neurons.

### Release Rate and Recycling pool of vesicles

The rate of vesicular release from depolarization induced by 90 mM High K+ Tyrode was significantly higher in syb2 overexpressing neurons than in WT **(Fig. 3B)**. We also detected a greater number of vesicles in the recycling pool when syb2 is overexpressed **(Fig. 3C)**. This suggests that overexpression of syb2 increases the vesicular release rate and the number of available functional synaptic vesicles.

**Figure 3:**
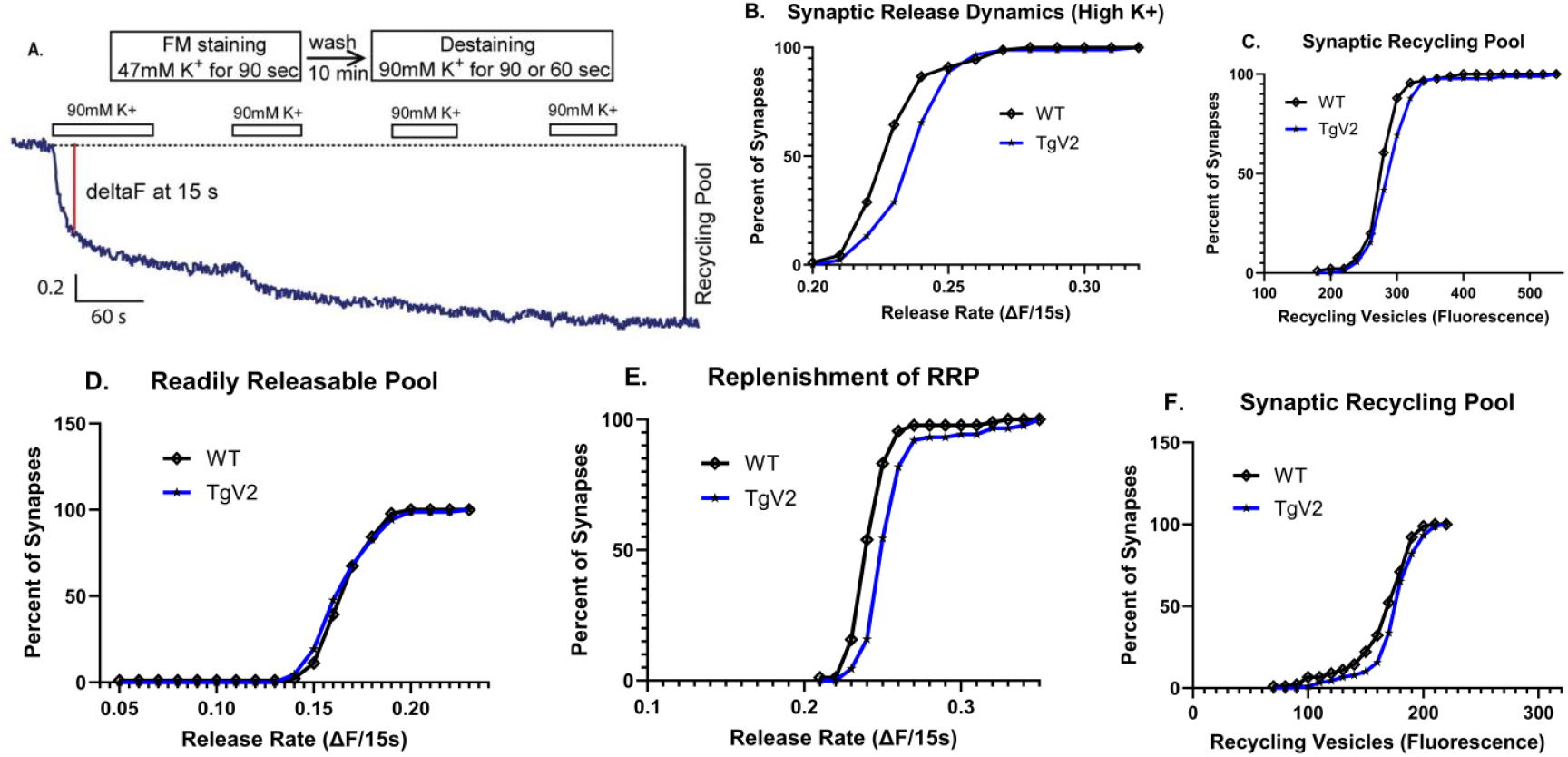
Live fluorescence imaging using High K+ protocol with FM1-43 Dye. **(A)**. Schematic of FM dye staining and destaining. **(B)** TgV2 synapses have a faster release rate than WT neurons (****p<0.0001; Kolmogorov–Smirnov test). **(C)** TgV2 neurons can release more synaptic vesicles, they have an increased recycling pool (**p=0.0012; Kolmogorov–Smirnov test). WT (n=9 brains; 2,118 synapses) and TgV2 (n=10 brains; 2,016 synapses). **Live fluorescence imaging using sucrose protocol with FM2-10 dye. (D)** Readily releasable synaptic pools in TgV2 and WT neurons WT (n=10 brains; 2,450 synapses) and TgV2 (n=12 brains; 2,104 synapses) p=0.3659; Kolmogorov-Smirnov test. **(E)** The rate of synaptic release in the second stimulation with depolarization (High K+) (****P<0.0001; Kolmogorov–Smirnov test). **(F)** TgV2 neurons can release more synaptic vesicles, they have an increased recycling pool (*p=0.0396; Kolmogorov–Smirnov test).

### Measuring readily releasable synaptic vesicular pool

In this experiment, the purpose of using hyperosmotic (500 mOsm) sucrose is to selectively release vesicles only from the readily releasable pool **(RRP)** of the synapse / synaptic vesicles **(Fig. 3D)**. Our data suggests that there is no difference in RRP between TgV2 and WT neurons. Importantly, after two-minute resting interval when we stimulated a second time using a High K+ Tyrode solution to depolarize the neurons, syb2 overexpressing neurons showed a significantly more vesicular release than WT **(Fig. 3E)**. This indicates that syb2 is a key regulatory factor for replenishment of vesicles, mobilizing vesicles from reserve pool to RRP.

#### Syb2 Overexpression Improves Neurotransmission in Cortical Neurons

Using whole cell patch clamp electrophysiology, we found that TgV2 primary cortical neurons had similar excitatory synaptic (EPSC) responses to field stimulation at low frequencies (-557.5 pA vs -495.7 pA, n=32-28, p= 0.2834; unpaired t-test). Paired Pulse Facilitation ratios showed no statistical difference between WT and TgV2 across the set range on time intervals from 10 to 500 ms **(Fig. 4B)**.

**Figure 4:**
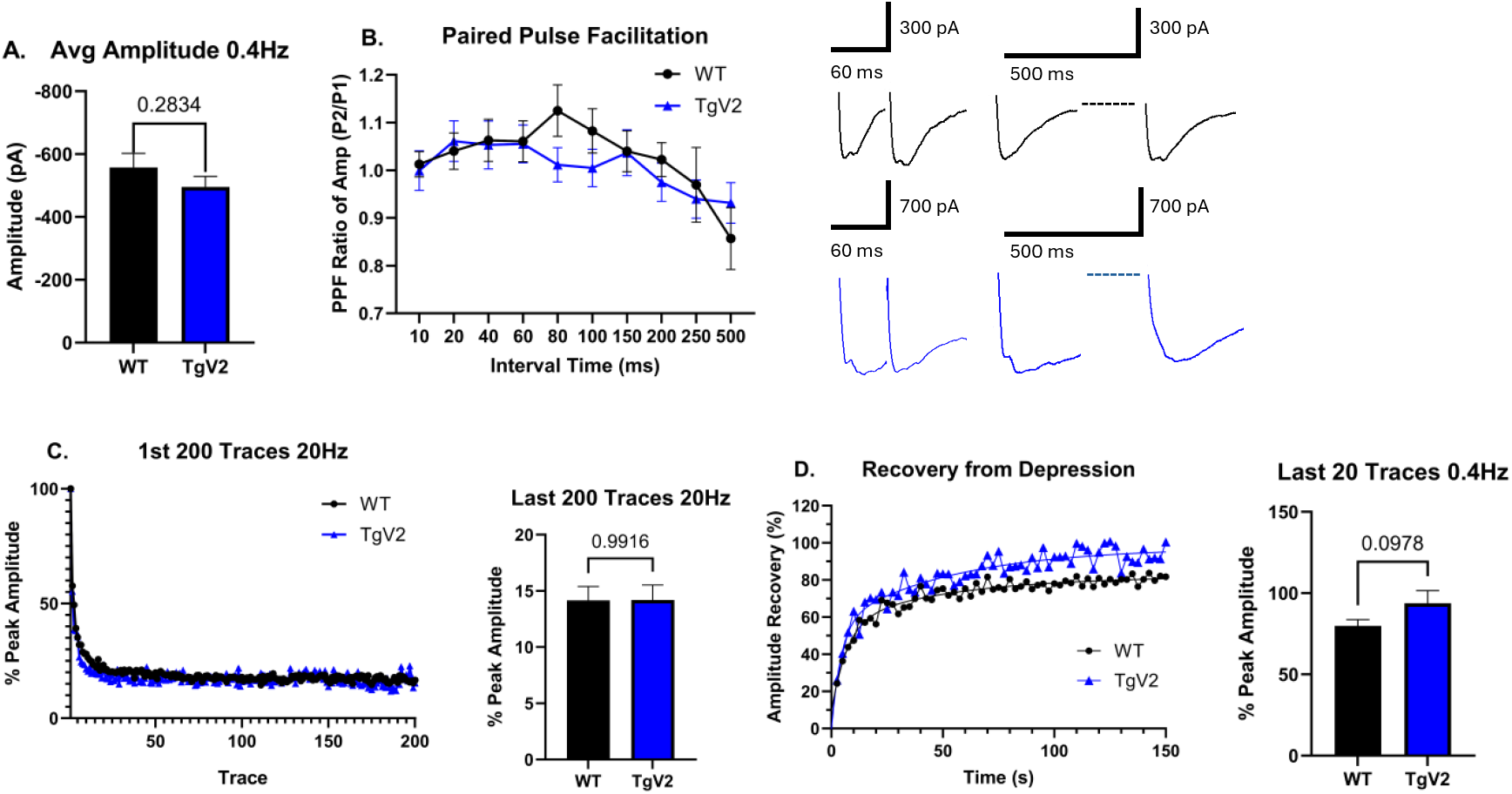
Whole-cell patch clamp on TgV2 neurons shows increased recovery of synaptic transmission compared to WT neurons (WT n=32 neurons) (TgV2 n=28 neurons). **(A)** Average amplitude of 1^st^ 0.4Hz **(B)** Percentage of peak EPSC amplitude during the last 200 traces at 20Hz showed no difference between genotypes. **(C)** 0.4Hz stimulation following 20Hz. TgV2 neurons show a greater percent recovery of peak amplitude during the last 20 traces of 0.4Hz (p=0.0978; unpaired t-test). **(D)** Paired Pulse Facilitation ratios showed no statistical different between WT and TgV2 across the set range on time intervals: 10ms, 20ms, 40ms, 60ms, 80ms, 100ms, 150ms, 200ms, 250ms, 500ms.

At 20 Hz stimulations TgV2 EPSCs depressed at a similar rate as WT neurons (percentage of WT peak amplitude = 14.17% vs percentage of TgV2 amplitude = 14.18% during last 200 traces of 20 Hz stimulation) **(Fig.4C)**. In the recovery period, during 60 stimuli at 0.4 Hz, TgV2 neurons also showed a significantly greater rate and percentage of EPSC recovery from depression following 20 Hz stimulation for one minute (average EPSC amplitude after 100 s during the last 20 traces at 0.4 Hz recovered to 79.84% in WT vs 93.82% in TgV2) **(Fig.4D)**. The recovery rate was most evidently faster at the beginning; it took only around 15 s for TgV2 to reach 63% recovery to the original EPSC amplitude compared to 20s for WT **(Fig.4D)**.

Recovery could be fitted with second order exponential equation with fast time constants of 4.3s for TgV2 and 6.8s for WT. Slow time constants were similar (52s and 51s, respectively) **(Fig.4D)**.

## Discussion

The key findings from our research indicated that maintaining expression levels of syb2 provides an effective preventative mechanism against cognitive decline during aging. To prove that syb2 levels are overexpressed in this model, we used a western blot assay using a monoclonal antibody for syb2 that recognizes both endogenous and syb2-eCFP fusion protein levels. It was found that this TgV2 mouse model maintains approximately twice the syb2 levels observed in severely aged WT hippocampus samples **(Fig.1B)**. This finding corresponds with the improved spatial memory and synaptic plasticity observed in severely aged TgV2 mice. This result also further confirmed our previously reported data that showed syb2 levels at or below 50% of that in young WT controls in 6-months old mice induced cognitive deficits ^23^. Using hippocampal slices from these aged animals, LTP recordings from TgV2 mice consistently indicated better synaptic plasticity in the CA1 Schaffer collaterals of TgV2 mice **(Fig. 2C)**. The improved ability of TgV2 neurons to maintain changes in synaptic strength within the aged hippocampus coincides with our MWM results that show severely aged TgV2 mice better retaining memory of the platform location one week after training **(Fig. 1F-J)**. This sustained plasticity during aging may not be relevant to object recognition as no differences in memory were observed between WT and TgV2 mice during novel object recognition tests **(data not shown)** Moreover, it was observed from input-output curves of these same slices that TgV2 mice have an extended range of potentiated responses to an increasing range of stimuli after high frequencies stimulation **(Fig.2D-E)**, which indicates that the overexpression of syb2 in TgV2 mice supports persistent synaptic weight changes after the induction of LTP. Further mechanistic support of this finding was obtained using patch-clamp electrophysiology and live-fluorescence imaging on primary cortical neurons. We found that vesicular release rates and the size of synaptic recycling pools were greater in TgV2 neurons compared to WT. TgV2 neurons displayed a faster vesicular release rate during depolarization using high KCl and FM1-43 dye **(Fig.3B)** or FM2-10 dye **(Fig.3E)**. These results imply that the role of syb2 overexpression is affecting the vesicular release from the recycling pool rather than the docked vesicles in the readily releasable pool as there were no differences when synaptic release was triggered by hyperosmolality using sucrose **(Fig.3D)**, while in both protocols TgV2 neurons had increased recycling pool of vesicles, as detected by a greater change in overall fluorescence **(Fig.3C, Fig.3F)**. Finally, to investigate the synaptic efficacy of TgV2 neurons we performed whole-cell patch clamp on primary cortical neurons and found that the average evoked excitatory post synaptic current (EPSC) was similar in WT and TgV2 overexpressing cortical neurons, which is in good agreement with unaltered RRP in the TgV2 neurons as seen with FM dye fluorescence imaging **(Fig. 3D)**. Although syb2 overexpression did not lead to better maintenance of EPSCs during 20 Hz **(Fig.4C)** or 10Hz stimulation **(data not shown)**, but it did improve the recovery of EPSC peak amplitude during the 0.4 Hz recovery period **(Fig.4D)**. This finding matches our FM imaging data and clearly indicates that TgV2 neurons have a significantly greater effect on the vesicular recycling pool compared to WT neurons. This, however, did not result in less synaptic fatigue or depression during high neuronal activity. We suggest that this may lead to higher probability of induction of synaptic plasticity, which is likely according to our CA1 LTP results.

It should be noted that, although the TgV2 line’s phenotype for prevention against deficiencies in spatial memory and plasticity were not observed until approximately 18-months old, it does not disqualify the results seen in primary cortical neurons isolated from neonatal mouse pups as causative mechanisms for this prevention. Syb2 levels in 6-8-month-old mice may be high enough to maintain proper cognition, while in the aged model the role of syb2 becomes more crucial for maintaining proper transmission as the levels of proteins within the SNARE complex decline.

Syb2 as biomarker of brain aging and neurodegeneration: The clinical relevance of this research is further highlighted by recent recognition of Syb2/VAMP2 as a biomarker of brain health, neurodegeneration and cognitive function. One may speculate that increased syb2 levels can lead to enhanced cognitive reserve, based on the observation of quicker synaptic responses and larger recycling vesicular pools in TgV2. Alternatively, the mechanism could involve also how excitatory synapses of TgV2 neurons recovered from depression – they were quicker than those in age matched WT controls. This would explain it was reported that higher syb2 mRNA levels may be associated with better cognition during aging ^23^.

Age is the most important risk factor for AD^26-32^. Importantly, the severity of cognitive impairment is not closely associated with senile plaque load, but best correlated with synapse loss, synaptic dysfunction and amount of fibrillary tangles^28, 33-37^. Syb2 levels were found to be lower in multiple brain regions of AD patients ^38^. Synaptobrevin was the only vesicle protein reduced in AD in all four brain regions, including occipital cortex and caudate nucleus ^39^. Interestingly, in Alzheimer’s disease loss of other synaptic proteins like synaptophysin, syntaxin, and SNAP-25 is a late-stage phenomenon associated with moderate to severe clinical grades of dementia and occurring only at pathological Braak5/6 stages with profound degeneration ^40^. VAMP2 levels is plasma are elevated if neurodegeneration destroys synapses. This was confirmed in both AD and PD patients ^41-44^. Syb2 is such a good reliable biomarker that recent clinical trials have used syb2 levels in CSF or plasma of AD patients as proof for treatment efficacy: if the drug protects from neurodegeneration, it reduces syb2 in CSF and plasma. For instance, sabirnetug treatment reduced CSF syb2 levels in the ongoing Intercept-AD clinical trial ^45^.

Beside the interaction with other SNARE proteins, Syb2 has a well-established physiological binding to alpha synuclein ^46^, which forms Lewy bodies in frontotemporal dementia (FTLD). Interestingly, VAMP2 levels are also increased in the CSF of patients with Lewy body dementia ^43^. Syb2 is emerging as a central hub in neurodegenerative disorders, as syb2 interacts with all major class of aggregated proteins in the most severe neurodegenerative disorders, like AD, FTLD and Parkinson’s disease, all leading to dementia.

### Limitations of this study

The results of our experiments are certainly generalizable to other mammalian species, however, there are three factors to be considered. 1. TgV2 is a transgenic mouse model and genomic integration of an artificial gene may cause unintended effects. Therefore, we identified the location of the transgene in the mouse genome and found five copies inserted in a genomic wasteland section of Chr. 15 ^25^. Thus, no major genomic interference with other genes is expected. The lack of significant behavioral difference between TgV2 and WT mice at 6–8-month-old affirms that no unintended effects of the transgene insertion happened. 2. The fusion of ECFP to syb2 might hinder function of syb2. This is unlikely as the design of TgV2 transgenic animal was based on our previous findings, showing a fully functional syb2 with this fluorescence tag attached via a flexible linker to the N-terminal of syb2 ^47^. 3. General aging effects of syb2: in theory one may argue that effect of syb2 may be more general than enhancing synaptic plasticity and may slow down the aging process. Our data neither supports nor refuses this hypothesis, which may require further experiments.

### Conclusions

To our best knowledge, this study is the first to investigate the effects of syb2 overexpression on age-related cognition in mice using the novel TgV2 transgenic mouse line. In this first functional analysis of the novel TgV2 mouse model our primary focus was on the role of syb2 in maintaining proper vesicular fusion and neurotransmitter release from the presynaptic membrane, and how increasing syb2 levels affects cognitive decline in aging. The findings reported demonstrate that the overexpression of syb2 has the potential to prevent memory deficits and synaptic dysfunction commonly reported in the neuropathology of the aged brain.

## Methods

### Animals

PI’s laboratory is approved to use this mouse model for better understanding synaptic neurotransmission. We have already obtained all mouse strains and successfully breed and maintain them in our colony.

### Generation of TgV2 mice

#### Vector Construction

First, pThy1-ECFP-C1 was made by cloning a 700-bp (XhoI-BsrGI) ECFP coding sequence into the Thy1 mini gene vector (generous gift of Dr. M. Goedert, MRC Laboratory Cambridge, UK) that contains the Thy1 genes without exon 3 and flanking introns to achieve neuron-specific expression ^48^. Second, an XhoI site was introduced by mutagenesis to the N-terminus of bovine synaptobrevin 2 (Syb2) in pCMV18-1a to generate pCMV18-1a-wp using the following oligonucleotides: 5’-CCG CTG CCA AGT CCT CGA GTC CGC TGG CCC CCG C and 5’-GCG GGG GCC AGC GGA CTC GAG GAC TTG GCA GCG G. Finally, a 1.5-kb XhoI-XhoI fragment from pCMV18-1a-wp was inserted into the XhoI site of pThy1-ECFP-C1 to generate the final transgenic vector pThy1-ECFP-Syb2.

#### Generation of Transgenic Mice

Transgenic mice were generated by injection of gel-purified pThy1-Syt1-ECFP DNA into fertilized oocytes using standard techniques ^49^. Transgenic founders were identified by DNA dot blotting and confirmed by PCR (see below). Eighteen founders from six injections were backcrossed to BL6SJL/F1 hybrid mice for 1–2 generations before protein and morphology analysis, and the two lines with the highest expression levels were retained for further experiments. The genotypes of transgenic mice were determined by PCR using the following primers, wh76 (TGG TGA ACC GCA TCG AGC TG, from ECFP) and wh79 (CGT TCA CCC TCA TGA TGT CC, from Syb2).

### Morris Water Maze

The Morris Water Maze (MWM) assay is commonly used to assess spatial learning and memory in rodents. Mice are trained in a pool to find a hidden platform over a series of trials throughout the weekdays and are then tested in a follow-up trial to gauge spatial learning memory abilities. The materials and methods included in our protocol consisted of the following. The pool used for this assay had a diameter of 100cm and a depth of 15cm. A white circular wall, containing four large images in the N, S, E, and W quadrants, was placed around the ends of the pool to block any possible external visual stimuli. The platform was 10cm in height and 6.5cm in height and the water level was kept 1-2 cm above the platform to allow the mice to enter easily. A constant water temperature of approximately 22°C was maintained through the experiment. Mice were trained for four days with four trials a day, sixty seconds maximum per trial if the platform was not found. After a gap period of one week, the platform was removed, and mice were allowed to swim for sixty seconds. The first cohort consisted of young adult mice (6-8 months old) and an aged cohort of approximately 2-years-old.

### Rotarod

Training consisted of three 2-minute trials at 10 revolutions per minute to habituate the mice and accustom them to balance on the Rotarod device (IITC Rotarod, California, USA). The testing phases were three 5-minute trials accelerating from 4-45 revolutions during each trial.

### Long-Term Potentiation

LTP was recorded on hippocampal slices (350µm) on a P5002A multi electrode array (MEA, AlphaMed, Japan), as described^50^. We stimulated the CA1 region with 30µA for 10 minutes at baseline responses before introducing 3×100Hz tetanic stimulation with 50µA. Immediately after, we returned to the baseline stimulus and recorded responses for 1 hour. We generated field excitatory post synaptic potentials (fEPSPs) on the Schaffer collateral pathway in the CA1 region by stimulating electrodes downstream in either CA3 or at a reasonable distance within CA1 itself **(Fig. 5A) at stimulus intensity evoking 30-40% of maximal fEPSP**. Tetanus was induced using three trains of 100 Hz stimuli for 1 s.

### Neonatal Primary Cortical Neuronal Cultures

Neonatal primary cortical neurons were isolated from mouse pups at P0 and were cultured for 12-14 days in vitro before using them for experimentation. The minimal essential medium used for neuronal cultures contained 5 g/L glucose, 0.1 g/L transferrin, 0.25 g/L insulin, 0.3 g/L glutamine, 5%–10% FBS, 2% B-27 supplement, and 1 μM cytosine arabinoside ^20, 25, 50^.

### Live Fluorescence Imaging

Primary cortical neurons were isolated from neonatal mouse pups and cultured on coverslips for 12-14 days. The live fluorescence imaging techniques shown in our preliminary data use two types of FM dye: FM2-10 for Sucrose protocol and FM1-43 for High K+ protocol. Olympus IX73 inverted microscope and Photometrics BSI 2048 CMOS camera were used for imaging. The fluorescence dye is added to a small aliquot of high concentration potassium (High K+) Tyrode solution and placed over the cover slip to induce vesicular release and fusion of the dye to the internal membrane of these vesicles. During the 90 second interval, these stained vesicles will undergo endocytosis into synapses and serve as a marker for synaptic release.

### Whole-Cell Patch Clamp

The patch clamp setup included an Olympus CK40 microscope, D1320 digitizer, Axopatch 200 amplifier and Burleigh 5300 micromanipulator. Pipette solution (mM): 10 NaCl, 110 Cs-methane-sulfonate, 20 TEA-Cl hydrate, 10 HEPES, 0.1 CaCl_2_ 2H_2_O, 4 Na_4_-BAPTA, 4 Mg-ATP, 0.3 Na_2_-GTP, 10 Lidocaine N-ethyl-bromide. 7.3 pH at 290mOsm.

100mM picrotoxin, a GABA-A receptor antagonist, was added to the bath solution to block inhibitory currents. Neurons were simulated at a set order of frequencies, including 0.4 Hz, 20 Hz, and 0.4 Hz, to acquire the EPSC peak amplitudes during baseline, depression, and recovery, respectfully. One minute after the 2^nd^ 0.4 Hz recovery period, paired pulse facilitation protocol was run using a gradually increasing time interval: 10ms, 20ms, 40ms, 60ms, 80ms, 100ms, 150ms, 200ms, 250ms, and 500ms.

## Acknowledgements

We thank Dr. Thomas C. Südhof for his support to develop the TgV2 mice.

## Authors contribution

Study concept and design: FD and JBM; drafting of the manuscript: FD and JBM; Western blots and behavioral studies: AMR, AS, and JBM. All other experiments: JBM analysis: acquisition, analysis, and interpretation: JBM and FD; reading and approval of the manuscript: all authors.

## Data availability

The data supporting this study’s findings will be available from the corresponding author upon reasonable request.

## Declarations

### Ethics approval and consent to participate

All animal experiments were approved by the IACUC of Augusta University.

### Consent for publication

Not applicable.

